# Is the sky the limit? On the expansion threshold of a species’ range

**DOI:** 10.1101/234377

**Authors:** Jitka Polechová

## Abstract

More than a hundred years after Grigg’s influential analysis of species’ borders, the causes of limits to species’ ranges still represent a puzzle that has never been understood with clarity. The topic has become especially important recently as many scientists have become interested in the potential for species’ ranges to shift in response to climate change – and yet, nearly all of those studies fail to recognise or incorporate evolutionary genetics in a way that relates to theoretical developments. I show that range margins can be understood based on just two measurable parameters: i) the fitness cost of dispersal – a measure of environmental heterogeneity – and ii) the strength of genetic drift, which reduces genetic diversity. Together, these two parameters define an *expansion threshold*: adaptation fails when genetic drift reduces genetic diversity below that required for adaptation to environmental heterogeneity. When the key parameters drop below this *expansion threshold* locally, a sharp range margin forms. When they drop below this threshold throughout the species’ range, adaptation collapses everywhere, resulting in either extinction, or formation of a fragmented meta-population. Because the effects of dispersal differ fundamentally with dimension, the second parameter – the strength of genetic drift – is qualitatively different compared to a linear habitat. In two-dimensional habitats, genetic drift becomes effectively independent of selection. It decreases with *neighbourhood size* – the number of individuals accessible by dispersal within one generation. Moreover, in contrast to earlier predictions, which neglected evolution of genetic variance and/or stochasticity in two dimensions, dispersal into small marginal populations aids adaptation. This is because the reduction of both genetic and demographic stochasticity has a stronger effect than the cost of dispersal through increased maladaptation. The *expansion threshold* thus provides a novel, theoretically justified and testable prediction for formation of the range margin and collapse of the species’ range.

**Author summary:** Gene flow across environments has conflicting effects: while it increases the genetic variation necessary for adaptation and counters the loss of genetic diversity due to genetic drift, it may also swamp adaptation to local conditions. This interplay is crucial for the dynamics of a species’ range expansion, which can thus be understood based on two dimensionless parameters: i) the fitness cost of dispersal – a measure of environmental heterogeneity – and ii) the strength of genetic drift – a measure of reduction of genetic diversity. Together, these two parameters define an *expansion threshold*: adaptation fails when the number of individuals accessible by dispersal within one generation is so small that genetic drift reduces genetic diversity below the level required for adaptation to environmental heterogeneity. This threshold provides a novel, theoretically justified and testable prediction for formation of a range margin and a collapse of a species’ range in two-dimensional habitats.

## Introduction

Species’ borders are not just determined by the limits of their ecological niche [1,2]. A species’ edge is typically sharper than would be implied by continuous change in the species’ environment (reviewed in [3, Table 2]). Moreover, although species’ ranges are inherently dynamic, it is puzzling that they typically expand rather slowly [4]. The usual – but tautological – explanation is that lack of genetic variation at the range margin prevents further expansion [5]. Indeed, a species’ range edge is often associated with lower neutral genetic variation [3,6–11], suggesting that adaptive genetic variation may be depleted as well [12]. Yet, why would selection for new variants near the edge of the range not increase adaptive genetic variance, thereby enabling it to continuously expand [5,13]? Haldane [14] proposed a general explanation: even if environmental conditions vary smoothly, “swamping” by gene flow from central to marginal habitats will cause more severe maladaptation in marginal habitats, further reducing their population density. This would lead to a sharp edge to a species’ range, even if genetic variance at the range margin is large. However, the consequences of dispersal and gene flow for evolution of a species’ range continue to be debated [15–18]: a number of studies suggest that intermediate dispersal may be optimal [19–23]. Gene flow across heterogeneous environments can be beneficial, because the increase of genetic variance allows the population to adapt in response to selection [13]. Current theory identifies that local population dynamics, dispersal, and evolution of niche-limiting traits (including their variance), and both genetic and demographic stochasticity are all important for species’ range dynamics [13,19–21,24–28]. Yet, these core aspects have not been incorporated into a single study that would provide testable predictions for range limits in two-dimensional habitats.

As Haldane [14] previously pointed out, it is important to consider population- and evolutionary dynamics across a species’ range jointly, as their effects interact. Due to maladaptation, both the carrying capacity of the habitat and the population growth rate are likely to decrease – such selection is called *hard* [31]. Classic deterministic theory [24] shows that when genetic variance is fixed, there are two stable regimes of adaptation to a spatially varying optimum (see Fig. 1): i) a *limited adaptation*, where a population is only adapted to a single optimum or becomes a patchy conglomerate of discrete phenotypes or ii) continuous or *uniform* adaptation, which is stable when the genetic variance, measured in terms of its cost in fitness (*standing genetic load*) is large relative to the maladaptation incurred by dispersal between environments (*dispersal load*). Under *uniform adaptation*, a species’ range gradually expands – a stable boundary only forms when the genetic variance is too small to allow continuous adaptation to the spatially variable environment and hence *limited adaptation* is stable.

**Figure 1.**
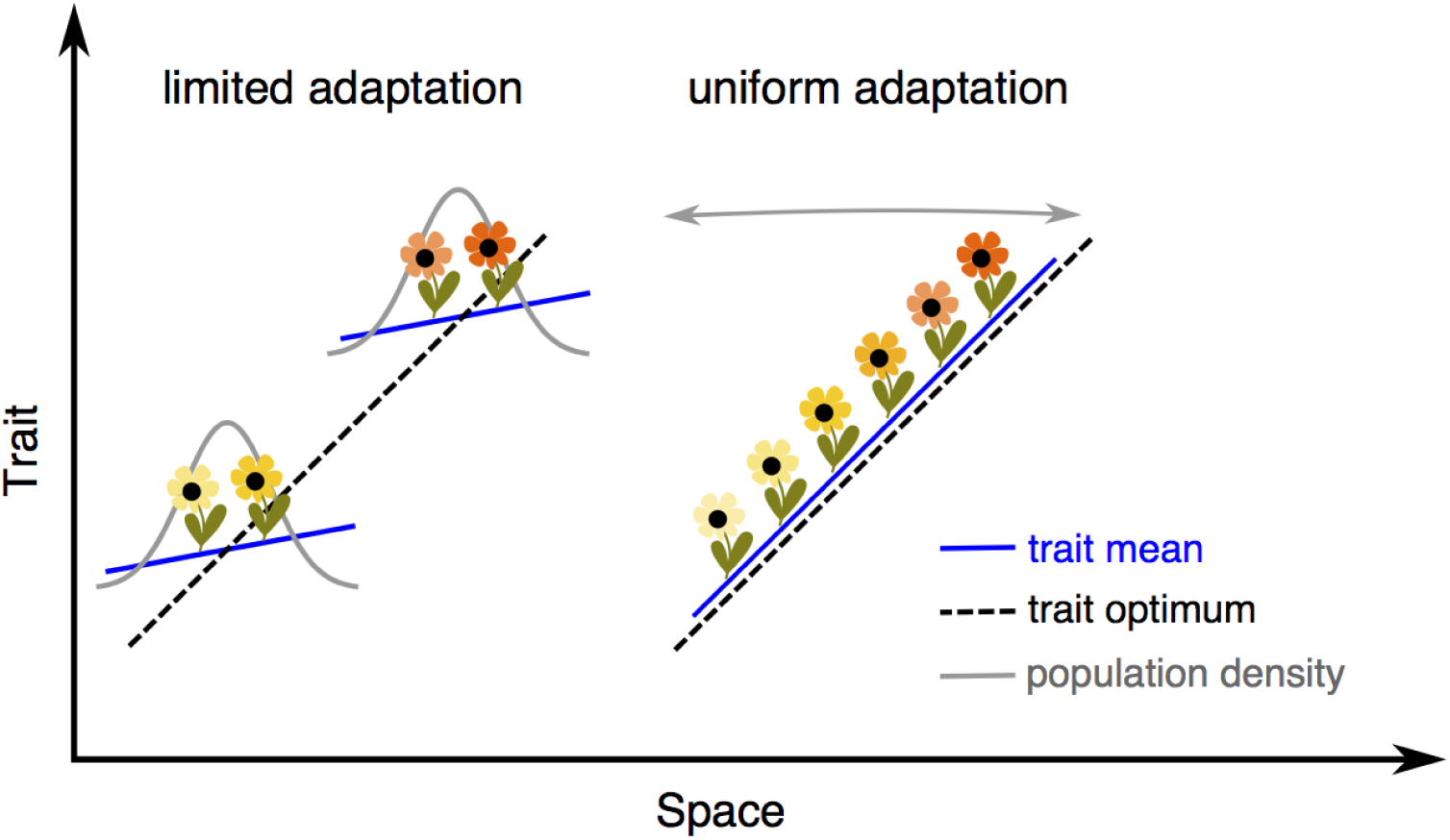
Two modes of adaptation. Assuming that genetic variance is fixed, deterministic theory [24] predicts that there are two modes of adaptation to an environmental gradient. When the effective environmental gradient *B* is steep relative to the genetic potential for adaptation *A*, clinal adaptation fails, and the population only matches the optimum at the very centre of its range (*limited adaptation*). These parameters can be understood as fitness *loads* scaled relative to the strength of density dependence *r*\* (see [24,29] and [30, Appendix D]). A is a measure of standing load due to genetic variance *Ar*\*, and *B* is a measure of dispersal load *B*^2^*r*\*^2^ – the maladaptation incurred by dispersal across heterogeneous environment. Thus conversely, when the standing load is large relative to the dispersal load, *A* > *B*^2^/2, a population adapts continuously, gradually expanding its range (*uniform adaptation*). Black dashed lines depict the trait optimum, blue lines depict the trait mean. Population density is shown in gray: it has a sharp and stable margin for *limited adaptation* but it is steadily expanding under *uniform adaptation*. Two sub-populations (or perhaps species) are given for illustration of *limited adaptation* – depending on further parameters and initial conditions (discussed in this study), a wide species’ range with *uniform adaptation* can collapse to a single population or fragment to multiple sub-populations.

When genetic variance can evolve, such a limit no longer exists in infinitely large populations: the population maintains continuous adaptation as the environmental gradient steepens [13]. Deterministic theory thus predicts that a sharp and stable boundary to a species’ range does not form when the environment changes smoothly. *Uniform* adaptation is the only stable regime when genetic variance can freely evolve in the absence of genetic drift [13], yet there is a limit to the steepness of the gradient. This limit arises because because both the *standing genetic load* and the *dispersal load* increase as the gradient steepens, reducing the mean fitness (growth rate) of the population: when the mean fitness approaches zero, the population becomes extinct. Obviously, ignoring genetic drift is then unrealistic. In finite populations, genetic drift reduces local genetic variance [32], potentially qualitatively changing the dynamics. Indeed, it has been shown that for one-dimensional habitats (such as rivers), a sharp range margin arises when the fitness cost of dispersal across environments becomes too large relative to the efficacy of selection versus genetic drift [26]. However, most species live in two-dimensional habitats. There, allele frequencies can fluctuate over a local scale as the correlations between them decline much faster across space than they do in linear habitats [33, Fig.3] and the effect of genetic drift changes qualitatively, becoming only weakly dependent on selection [34]). Is there still an intrinsic threshold to range expansion in finite populations when dispersal and gene flow occur over two-dimensional space, rather than along a line? If so, what is its biological interpretation?

## Results

I study the problem of intrinsic limits to adaptation in a two dimensional habitat. Throughout, I assume that the species’ niche is limited by stabilising selection on a composite phenotypic trait. This optimum varies across one dimension of the two-dimensional habitat – such as temperature and humidity with altitude. Demography and evolution are considered together. Selection is *hard*: both the rate of density-dependent population growth and the attainable equilibrium density decrease with increasing maladaptation. Both trait mean and genetic variance can freely evolve *via* change in allele frequencies, and the associations among them (linkage disequilibria). The populations are finite and both genetic and demographic stochasticity are included. The model is first outlined at a population level, in terms of coupled stochastic differential equations. While it is not possible to obtain analytical solutions to this model, this formalisation allows us to identify the effective dimensionless parameters which describe the dynamics. Next, individual based simulations are used to determine the driving relationship between the key parameters, and test its robustness. The details are described in Methods: Model.

The dynamics of the evolution of a species’ range, as formalised by this model, are well described by three dimensionless parameters, which give a full description of the system. Details are given in the Methods: Rescaling. The first dimensionless parameter carries over from the phenotypic model [24]: the effective environmental gradient *B* measures the steepness of the environmental gradient in terms of maladaptation incurred by dispersal across a heterogeneous environment. The second parameter is the neighbourhood size of the population, 𝒩, which can be understood as the number of diploid individuals within one generation’s dispersal range. Originally, neighbourhood size was defined by Wright [35] as the size of the single panmictic diploid population which would give the same probability of identity by descent in the previous generation. The inverse of neighbourhood size 1/𝒩 hence describes the local increase of homozygosity due to genetic drift. The third dimensionless parameter is the ratio *s/r*\* of the strength of selection *s* per locus relative to the strength of density dependence, *r*\*. Detailed description of the parameters can be found in Methods: Table 1.

In order to see how the rescaled parameters capture the evolution of a species’ range, I simulated 780 evolving populations, each based on different parameterizations, adapting to a linear gradient in the optimum. Depending on the parameters, the population either expands, gradually extending its phenotypic range by consecutive sweeps of loci advantageous at the edges, or the species’ range contracts or disintegrates as adaptation fails. Fig. 2 shows the results of the projection from a 10-dimensional parameter space of the individual-based model (see Methods: Individual-based Model and Table 1) into a two-dimensional space. The axes of Fig. 2 represent the first two compound dimensionless parameters: i) the effective environmental gradient *B* and ii) the inverse of neighbourhood size 1/𝒩 which describes the effect of genetic drift on the allele frequencies.

**Figure 2.**
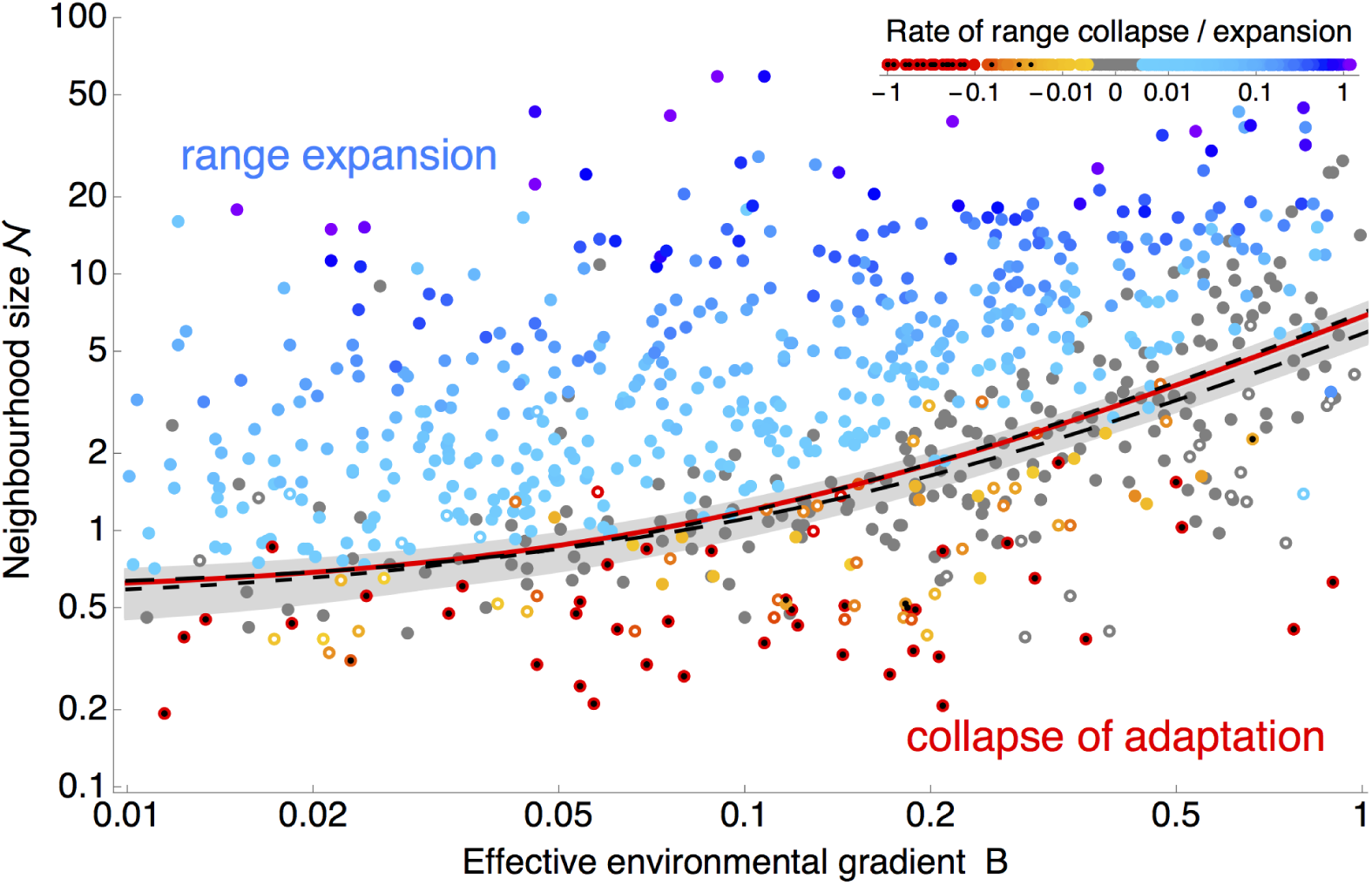
Two dimensionless parameters, the neighbourhood size 𝒩 and the effective environmental gradient *B* give a clear prediction whether a species’ range can expand (blue hues). The red line shows the fitted boundary between expanding populations (in blue) and collapsing ranges (red hues): populations expand above the *expansion threshold*, when 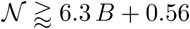. The grey region gives 95% bootstrap confidence intervals, whilst the dashed lines depict the predicted *expansion threshold* for weak selection, *s/r*\* < 0.005 (- -) and for strong selection, *s/r\** > 0.005 (– –). Stagnant populations, changing by less than five demes per 1000 generations, are shown in grey. Open circles denote populations where continuous adaptation has collapsed and the population consists of many discrete phenotypes adapted to a single optimum each (*limited adaptation*, Fig. 4), whilst local genetic variance is very small (defined by mean heterozygosity smaller than 10% of the predicted value in the absence of genetic drift). Simulations were run for 5000 generations, starting from a population adapted to a linearly changing optimum in the central part of the available habitat. Populations that went extinct are marked with a black dot. Note that both axes are on a log scale. The top corner legend gives the colour coding for the rate of range collapse and expansion in units of demes per generation; rates of collapse are capped at –1. The *expansion threshold* is fitted as a step function changing linearly along *B*: all blue dots are assigned a value of 1; all red dots and open circles are assigned a value of 0. The *expansion threshold* has a coefficient of determination *R*^2^ = 0.94, calculated from 589 simulations (all but well-adapted stagnant populations).

These two dimensionless parameters *B* and 𝒩 give a clear separation between expanding populations when the neighbourhood size 𝒩 is large relative to the effective environmental gradient *B* (shown in blue, Fig. 2) and the rest, where adaptation is failing. The separation gives an *expansion threshold*, estimated at 𝒩 ≈ 6.3 *B* + 0.56 (red line). Above the *expansion threshold*, populations are predicted to expand (see Fig. 3); below it, adaptation fails abruptly. If conditions change uniformly across space (as in these simulation runs, with constant gradient and carrying capacity), this means that adaptation fails everywhere – a species’ range then either collapses from the margins (Fig. 2, red hues) and/or disintegrates (Fig. 2, open circles), forming a fragmented metapopulation (i.e., a spatially structured population consisting of discrete locally adapted sub-populations with limited dispersal among them).

**Figure 3.**
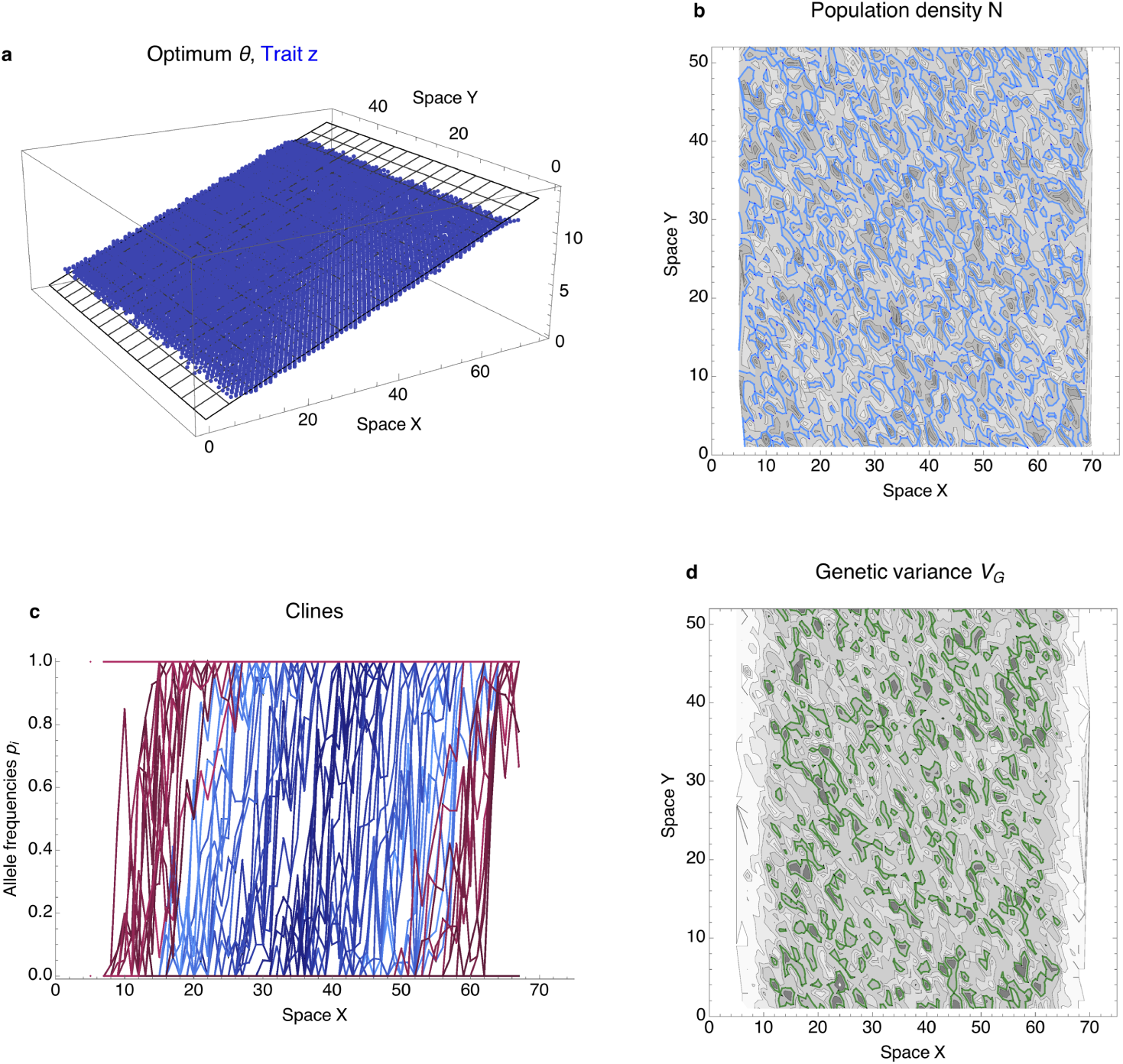
Uniform adaptation: above the *expansion threshold*, the population expands gradually through the available habitat. (a) Trait (in blue) closely matches the environmental gradient (grey) along the X axis. (b) Population steadily expands, whilst population density stays continuous across space, with 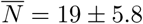 (mean ± standard deviation). The prediction at 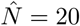 is shown by the blue contours; darker shading represents higher density. (c) Adaptation to the environmental gradient is maintained by a series of staggered clines: as each allele frequency changes from 0 to 1, the trait value increases by *α*. Population starts from the centre (blue hues reflect initial cline position relative to the centre of the range) and as it expands new clines arising from loci previously fixed to 0 or 1 contribute to the adaptation (in red hues). At each location, multiple clines contribute to the trait (and variance); clines are shown at Y = 25. (d) Genetic variance changes continuously across space with mean 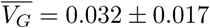, and stays slightly lower than is the deterministic prediction (green contours, *V_G_* = 0.045; higher variance is illustrated by darker shading). Deterministic predictions are based on [13], and are explained in Methods, along with the specification of the unscaled parameters. The population evolves for 2000 generations, starting from a population adapted to the central habitat. The predicted neighbourhood size is 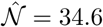, effective environmental gradient is *B* = 0.48.

When metapopulation forms, it exhibits an extinction and colonisation dynamics. The sub-populations drift freely around the neutral spatial axis (keeping the trait optimum) and they also drift along the environmental gradient, where the location changes together with the sub-population’s trait mean. Over time, the metapopulation very slowly collapses to a single trait value. The sub-populations forming this metapopulation have only a very narrow phenotypic range and maintain locally only minimal adaptive variance. They correspond to the *limited adaptation* regime identified for a phenotypic model with genetic variance as a parameter [24]. In contrast to one-dimensional habitats [26], these patchy metapopulations are stabilised by dispersal from surrounding subpopulations in the two-dimensional habitat and can thus persist for a long time. An example of such a metapopulation is given in Fig. 4.

**Figure 4.**
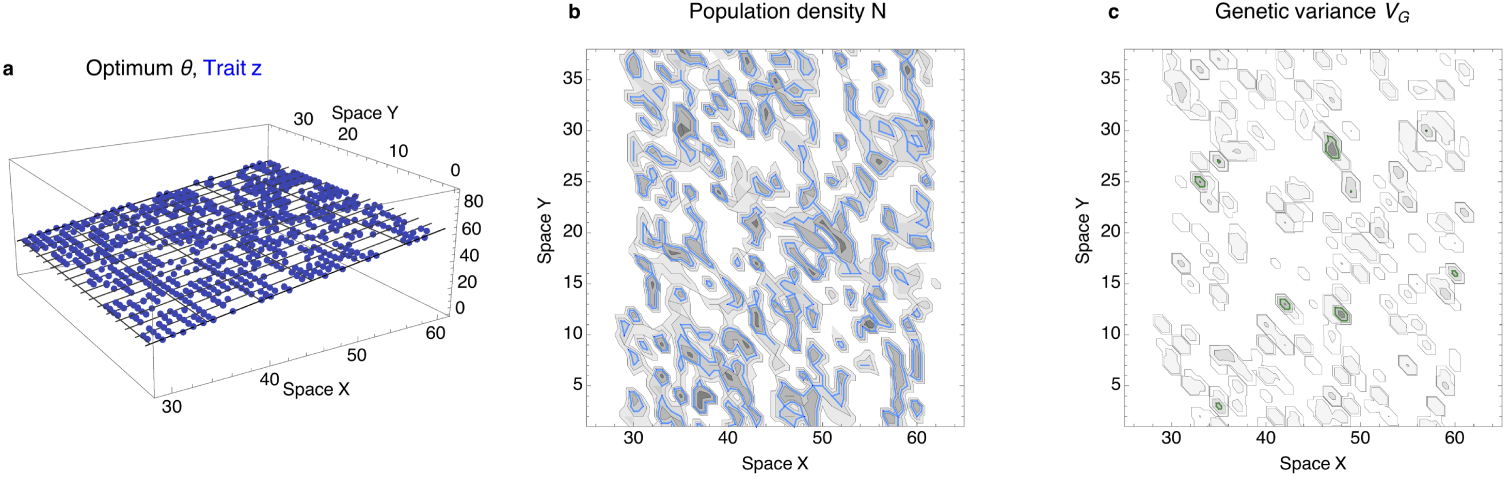
A metapopulation can form when the population is below the *expansion threshold* throughout its range. The population fragments rapidly (within tens of generations) to small patches of tens to a few hundred of individuals, whilst losing local adaptive variation. In two-dimensional habitats, such a metapopulation with *limited adaptation* can persist for long times. Nevertheless, the population very slowly contracts, eventually forming a narrow band adapted to a single optimum. (a) The distribution of phenotypes across space is fragmented. (b) The sub-populations are transient, although they are stabilised by dispersal across space, especially along the neutral direction with no change in the optimum (Y). Locally, the population density may be higher than under *uniform adaptation*; blue contours depict the deterministic prediction for population density under *uniform adaptation, N* = 3. The realized density is about 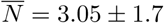 (standard deviation); darker shading represents higher density. (c) The adaptive genetic variance is low on average 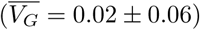 – about an order of magnitude lower than would be maintained by gene flow under *uniform adaptation* (shown in green contours, *V_G_* = 0.23). Typically, only a few clines in allele frequencies contribute to the genetic variance within a sub-population. The parameterization and predictions are detailed in the Methods: Individual-based model; predicted neighbourhood size is 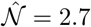, effective environmental gradient is *B* = 0.48. Shown here after 5000 generations – the population collapses to a narrow band (at *X* = 45) after a further 20.000 generations and then appears persistent.

Interestingly, the third dimensionless parameter *s/r*\* has no detectable effect on the form of the expansion threshold. In other words, whilst the *expansion threshold* reflects the total fitness cost of dispersal in a heterogeneous environment, it appears independent of the strength of selection per locus s: the dashed lines in Fig. 2 compare the estimated expansion threshold for small and large *s/r*\*. Increasing the strength of selection is inefficient in aiding drift-limited adaptation, in line with the expectation that the effect of genetic drift is only very weakly dependent on selection in two-dimensional habitats [27; see also Fig. S1]. This suggests that genetic basic of adaptation is not important for a drift-induced limit to a species’ range. Yet, it is plausible that there is another limit, where selection per locus becomes important [27], which arises when the optimum changes abruptly and even when the population (neighbourhood) size is large (i.e., in an entirely different regime). A dedicated synthesis connecting the step-limited and drift-limited regimes would be of a clear interest. Importantly, once genetic drift starts to have an effect, the habitat needs to be fairly broad to be two-dimensional [36]. In narrow habitats (such as in [27]), residual dependency of drift-induced *expansion threshold* would be expected [26]. Note that the apparent independence of the *expansion threshold* on *s/r*\* does not imply that rate of range expansion should be also independent of the strength of selection.

In nature, conditions are unlikely to change uniformly. Abiotic environment (such as temperature, precipitation, solar radiation) does not in general change in a linear and concordant manner [37], and neither does the biotic environment, such as the pressure from competitors and predators, which affects the attainable population density and can increase the asymmetry in gene flow [38, 39]. I now investigate whether adaptation fails near the *expansion threshold* as conditions change across space. For example, we can imagine that the population starts well adapted in the central part of the available habitat, and as it expands, conditions become progressively more challenging (see Fig. S2a); i.e. the effective environmental gradient *B* gets steeper. As the expanding population approaches the *expansion threshold*, adaptive genetic variance progressively decreases below the predicted value [13], which would be maintained by gene flow in the absence of genetic drift (Fig. 5a, grey dashed line). This is a result of an increased frequency of demes under *limited adaptation*, leading to higher rates of extinctions and re-colonizations, which reduce both adaptive and neutral diversity (see Fig. 5b). Range expansion then ceases at the *expansion threshold* as the genetic variance drops to the critical value where only *limited adaptation* is stable [24] assuming genetic variance is fixed (Fig. 5a, dotted line). This is because although populations can persist with *limited adaptation* (Fig. 4), the transient amount of genetic variance maintained under *limited adaptation* is almost never consistent with range expansion (see Fig. 2, open circles). On a steepening gradient, a sharp and stable range margin forms. This contrasts to uniformly changing conditions (linear gradient, constant carrying capacity), where populations steadily expand or contract.

**Figure 5.**
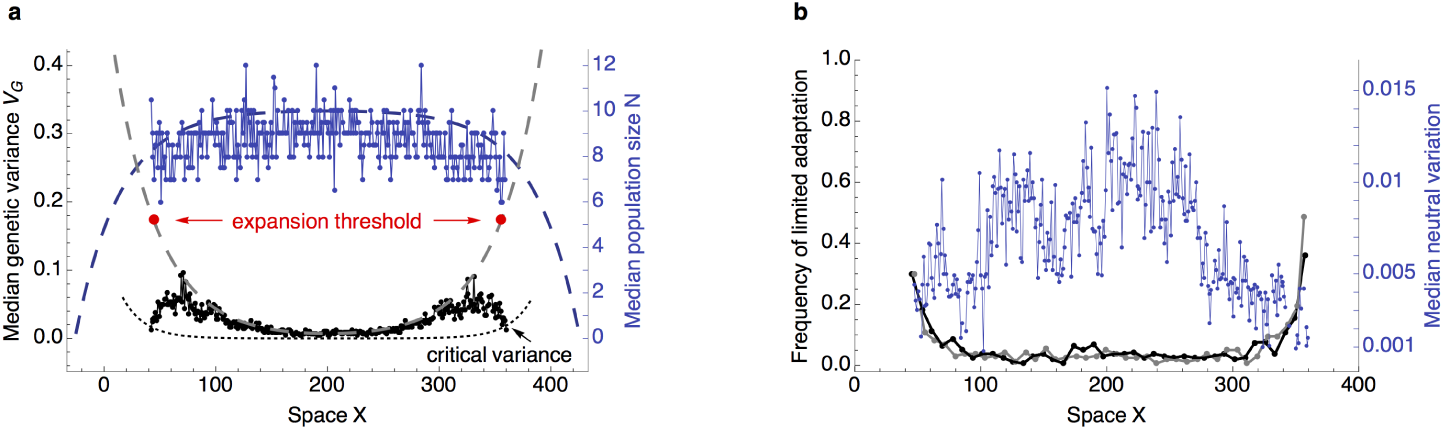
On a steepening environmental gradient, a sharp and stable range margin forms near the *expansion threshold*. This illustrative run shows that as the effective environmental gradient steepens away from the central location, adaptive genetic variance must increase correspondingly in order to maintain *uniform adaptation*. (a) Median population density stays fairly constant across the range (blue dots), following the deterministic prediction (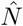, blue dashed line). Genetic variance (black dots) increases due to gene flow across the phenotypic gradient – the deterministic expectation is given by the gray dashed line. (See Methods: Model for details.) Yet as the environmental gradient steepens, genetic variance fails to increase fast enough and near the *expansion threshold*, adaptation fails. The dotted line gives the corresponding critical genetic variance, below which only *limited adaptation* is expected in a phenotypic model with a fixed genetic variance (*A* ≈ *B*^2^/2, where A is the standing genetic load; [24]). (b) As the environmental gradient steepens, the frequency of *limited adaptation* within the metapopulation increases (black and gray) and hence neutral variation decreases (blue). The black line gives the proportion of demes with *limited adaptation* after 50 000 generations, when the range margin appears stable; grey gives the proportion after 40 000 generations (depicted is an average over a sliding window of 15 demes). The median is given over the neutral spatial axis Y (with constant optimum); the trait mean, the population trait mean, variance and population density in two-dimensional space is shown in Fig. S2 which also lists all the parameters.

In a large population, the ability to adapt to heterogeneous environments is independent of dispersal: this is because both the local genetic variance (measured by *standing genetic load*), which enables adaptation to spatially variable environments, and the perceived steepness of the environmental gradient (measured by *dispersal load*), increase at the same rate with gene flow [13]. Yet, in small populations, dispersal is beneficial because the drift-reducing effect of dispersal overpowers its maladaptive effect. This is demonstrated in Fig. 6 – the neighbourhood size 𝒩 increases faster with dispersal than the effect of swamping by gene flow (*B*) does – hence, as dispersal increases, the population gets above the *expansion threshold*, where *uniform adaptation* can be sustained. Around the *expansion threshold*, a small change in dispersal (connectivity) can have an abrupt effect on adaptation across a species’ range and the species’ persistence. A small increase in dispersal can lead to recovery of *uniform adaptation* with an arbitrarily wide continuous range. Further increase of dispersal is merely enhancing the rate of range expansion at the expense of a slight cost to the mean fitness due to rising *dispersal load* and *standing load*, and can be associated with further costs, such as Allee effect (see eg. [17]). Therefore, the *expansion threshold* provides an interpretation for optimality of an “intermediate” dispersal, benefiting the species’ persistence.

**Figure 6.**
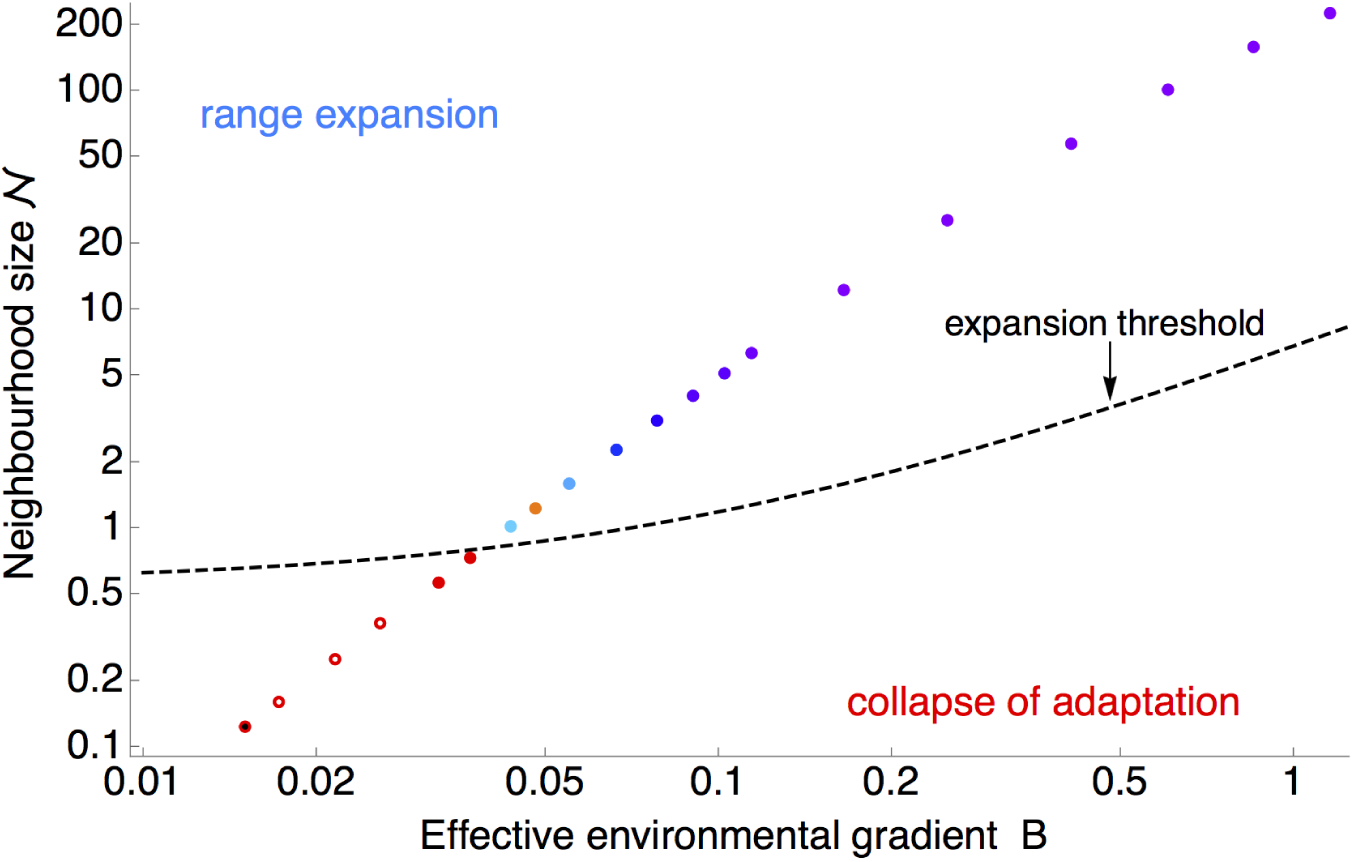
Dispersal aids adaptation in small populations because the neighbourhood size 𝒩 increases with the square of generational dispersal, whereas the effective environmental gradient *B* increases only linearly. This chart shows a set of simulated populations, with dispersal increasing from left to right and bottom to top. The hue of the dots indicates the rate of expansion (light to dark blue and purple) or collapse (orange to red). The rates of expansion and collapse are shown in dependency on *B* and 𝒩. Open circles indicate *limited adaptation*, where species’ range is fragmented and each subpopulation is only matching a single optimum, whilst its genetic variance is very small. As dispersal increases, population characteristics get above the *expansion threshold* (dashed line), and hence *uniform adaptation* becomes stable throughout the species’ range. Local population density stays fairly constant, around *N* = 3.5, whilst total population size increases abruptly above the *expansion threshold* as the population maintains a wide range (not shown). Parameters for these simulations are given in the Methods: Individual-based model; the scaling of 𝒩 and *B* with dispersal *σ* is clear from Table 1. The rate of range change is not significantly different from zero for the first three simulations above the *expansion threshold*; black centre (bottom left) indicates extinction.

## Discussion

Here I have shown that adaptation fails when positive feedback between genetic drift, maladaptation and population size reduces adaptive genetic variance to levels which are incompatible with continuous adaptation. The revealed *expansion threshold* differs qualitatively from the limit to adaptation identified previously [26] for a population living along a one-dimensional habitat. This is because in two dimensions, dispersal mitigates the loss of diversity due to genetic drift more effectively, such that it becomes (almost) independent of selection [34]. The *expansion threshold* implies that populations with very small neighbourhood sizes 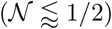, which suffer a severe reduction in neutral heterozygosity, will be prone to collapse based on demographic stochasticity alone. However, even in the absence of demographic stochasticity, genetic drift reduces the adaptive genetic variance required to sustain adaptation to a heterogeneous environment. The *expansion threshold* describes when this reduction due to genetic drift is incompatible with continuous adaptation, predicting a collapse of a species’ range. If the *expansion threshold* is reached as the species expands through its habitat, a sharp and stable range margin forms. If there is a drop below the *expansion threshold* throughout the species’ range, as after a sudden drop in carrying capacity, adaptation abruptly collapses throughout a species’ range. The result is either extinction, or a fragmented metapopulation consisting of a conglomerate of sub-populations, each adapted to a single phenotypic optimum. It follows that near a range margin, we expect increased range fragmentation, and a decrease in adaptive genetic variance. The threshold gives a theoretical base to the controversial issue of the importance of evolution (genetics) and ecology (demography) for assessing vulnerability of a species [40,41]. The predicted sharp species’ range edge is in agreement with the reported lack of evidence for *abundant centre* of a species’ range, which although commonly assumed in macroecological theory, has little support in data [3,11,42,43]. Lack of abundant centre is consistent both with uniform adaptation and with limited adaptation in a metapopulation.

The *expansion threshold* provides a general foundation to species-specific eco-evolutionary models of range dynamics [44]. Its components can be measured in wild populations, allowing to test the robustness of the theory. First, the effective environmental gradient *B* can be measured as fitness loss associated with transplant experiments on a local scale, relative to a distance of generational dispersal along an environmental gradient. The environmental gradient can include both biotic and abiotic effects, and their interactions [45] – notably, the effective environmental gradient *B* steepens due to increased asymmetry in gene flow when carrying capacity varies across space, e.g. due to partial overlap with competitors [39]. Second, the neighbourhood size 𝒩 can be estimated from neutral allele frequencies [46,47]. Estimates of neighbourhood size are fairly robust to the distribution of dispersal distances [48]. Though near the *expansion threshold*, both the noisiness of the statistics and the homozygosity will increase due to local extinctions and recolonizations [49]. An alternative estimate of neighbourhood size can be also obtained from mark-recapture studies, by measuring population density and dispersal (as an approximation for gene flow) independently [46].

Because the *expansion threshold* is free of any locus- or trait-specific measure, the result appears independent of genetic architecture, readily extending to multiple traits regardless of their correlations (c.f. [50–54]) – yet, the mean fitness will decline due to *drift load* as the number of traits independently optimised by selection increases [55, 56]. Hence if the fitness landscape is highly complex, the *expansion threshold* constitutes a lower limit. Naturally there can be further costs arising in a natural population which I have neglected here – such as the Allee effect [17]. In general, while the numerical constants may change when natural populations deviate in their biology from our model assumptions, the scale-free parameters identified in this study remain core drivers of the intrinsic dynamics within a species’ range. Notably, the early classic studies assuming fixed genetic variance [24] predicted that dispersal into peripheral populations is detrimental because it only inflates the effective environmental gradient *B*. Yet, when genetic variance can evolve, dispersal into small marginal populations also aids adaptation by increasing local genetic variance and by countering genetic drift. The net effect of dispersal into small marginal populations (below the *expansion threshold*) is then beneficial because their neighbourhood size increases faster with dispersal than the effective environmental gradient *B* steepens.

## Methods

### Model

I model evolution of a species’ range in a two-dimensional habitat, where both population dynamics and evolution (in many additive loci) are considered jointly. The coupling is *via* the mean fitness 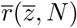, which gives the growth rate of the population, and decreases with increasing maladaptation: 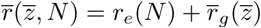. The ecological component of growth rate, *r_e_*, can take various forms: here the regulation is logistic, so that fitness declines linearly with density *N*: *r_e_* = *r_m_*(1 — *N/K*), where *r_m_* is the maximum per capita growth rate in the limit of the local population density *N* → 0. The carrying capacity *K* (for a perfectly adapted phenotype) is assumed uniform across space. The second term, 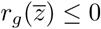, is the reduction in growth rate due to deviation from the optimum. Selection is stabilising: the optimum *θ* changes smoothly with one spatial dimension (*x*): for any individual, the drop in fitness due to maladaptation is *r_g_*(*z*) = − (*z* − *θ)*^2^/(2*V_s_*). Here *V_s_* gives the width of stabilising selection; strength of stabilising selection is *γ* = − *V_P_*/(2*V_s_*), where *V_P_* = *V_G_* + *V_E_* is the phenotypic variance. A population with mean phenotype 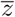 has its fitness reduced by 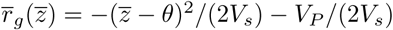. The phenotype *z* is determined by many di-allelic loci with allelic effects *α_i_*; the model is haploid, hence 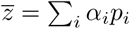, where *p_i_* is the allele frequency at locus *i*. Phenotypic variance is *V_P_* = *V_G_* + *V_E_*. The loss of fitness due to environmental variance *V_E_* can be included in 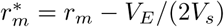; *V_E_* is a redundant parameter. Selection is *hard*: both the mean fitness (growth rate) and the attainable equilibrium density 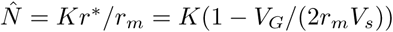 decrease with maladaptation. Expected genetic variance maintained by gene flow in the absence of genetic drift is *V_G_* = *bσV_s_* [13]; the contribution due to mutation is small, at mutation-section balance 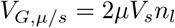, where *μ* gives the mutation rate per locus and *n_l_* the number of loci.

### Individual-based simulations

Discrete-time individual based simulations are set to correspond to the model with continuous time and space. The space is a two-dimensional lattice with spacing between demes of *δx* = 1. Every generation, each individual mates with a partner drawn from the same deme, with probability proportional to its fitness, to produce a number of offspring drawn from a Poisson distribution with mean of *Exp*[*r*(*z, N*)] (this includes zero). The effective diploid population density *N_e_* hence equals half of the haploid population density *N*, and 𝒩 = 4*πN_e_σ^2^* = 2*πNσ*^2^. The life-cycle is selection → mutation → recombination → birth → migration. Generations are non-overlapping and selfing is allowed at no cost. The genome is haploid with unlinked loci (the probability of recombination between any two loci is 1/2). The allelic effects *α_i_* of the loci combine in an additive fashion; the allelic effects are uniform throughout this study, *α_i_* = *α*. Mutation is set to *μ* = 10^−6^, independently of the number of loci. Migration is diffusive with a Gaussian dispersal kernel. The tails of the dispersal kernel need to be truncated: truncation is set to two standard deviations of the dispersal kernel throughout, and dispersal probabilities and variance are adjusted so that the discretised dispersal kernel sums to 1 [57, p. 1209]. Simulations were run at the computer cluster of IST Austria using *Mathematica 9 (Wolfram)*; the code used for simulations is available at request.

#### Parameters

There are in total 10 parameters in the individual-based model but only 7 are used to describe the model dynamics in continuous time. These are listed at the bottom of Table 1. They are the environmental gradient *b* = [0.012, 2], dispersal distance *σ* = [0.1,1.3], carrying capacity for a well adapted phenotype *K* = [3, 31], width of stabilising selection *V_s_* = [0.005, 6], the maximum intrinsic rate of increase *r_m_* = [0.2, 2] and the mutation rate *μ*, fixed to *μ* ≡ 10^−6^. The [*x,y*] interval gives the parameter range used in the 780 randomly sampled runs, with their distributions described in Fig. S3. The number of genes and demes are not included in the continuous time description (and hence the rescaling) because it assumes that space is not limiting, and that all loci have equivalent effect with no statistical associations among them. In the individual-based model, the habitat width is set to be wide enough to be effectively two-dimensional under diffusive dispersal for thousands of generations [36] – 100 dispersal distances *σ* along the neutral direction, and at least 10 cline (deterministic) widths wide along the gradient. The number of genes contributing to the adaptation across the species’ range is *n_l_* = [5, 2996], with the estimated number of locally polymorphic genes between 1 and 299. Since mutation rate is fixed at *μ* = 10^−6^, the genomic mutation rate has a wide range, *U* = [5 · 10^−6^, 3 · 10^−3^], with median of *U* = 10^−4^.

Parameters for Fig. 3 are: *b* = 0.18, *σ* = 0.52, *V_s_* = 0.23, *K* = 26.7, 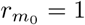 and *α* = 0.14, *s* = 0.04, *μ* = 10^−6^, 97 genes. Median genetic variance is at *V_G_* = 0.031, deterministic prediction 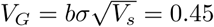 [13]. In Fig. 4, the parameters are: *b* =1, *σ* = 0.4, *V_s_* = 0.4, *K* = 4, *r_m_* = 1.2 and *α* = 0.1, *s* = 0.015, *μ* = 10^−6^, 874 genes. Median genetic variance within patches is around 0.02, whilst the maximum contribution by a single cline 1/4*α*^2^ = 0.0026; in contrast, variance maintained by gene flow under *uniform adaptation* [13] would be 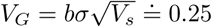. Parameters for Fig. 6 are *b*_0_ = 0.3, *σ* = [0.05, 3], *V_s_* = 1, *K* = 4, 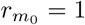 and *α* = 0.05, *s* = 0.1, *μ* = 10^−6^, 1000 genes, 1000 demes along X, 200 demes along Y. These populations evolved for 500 generations.

**Table 1.**
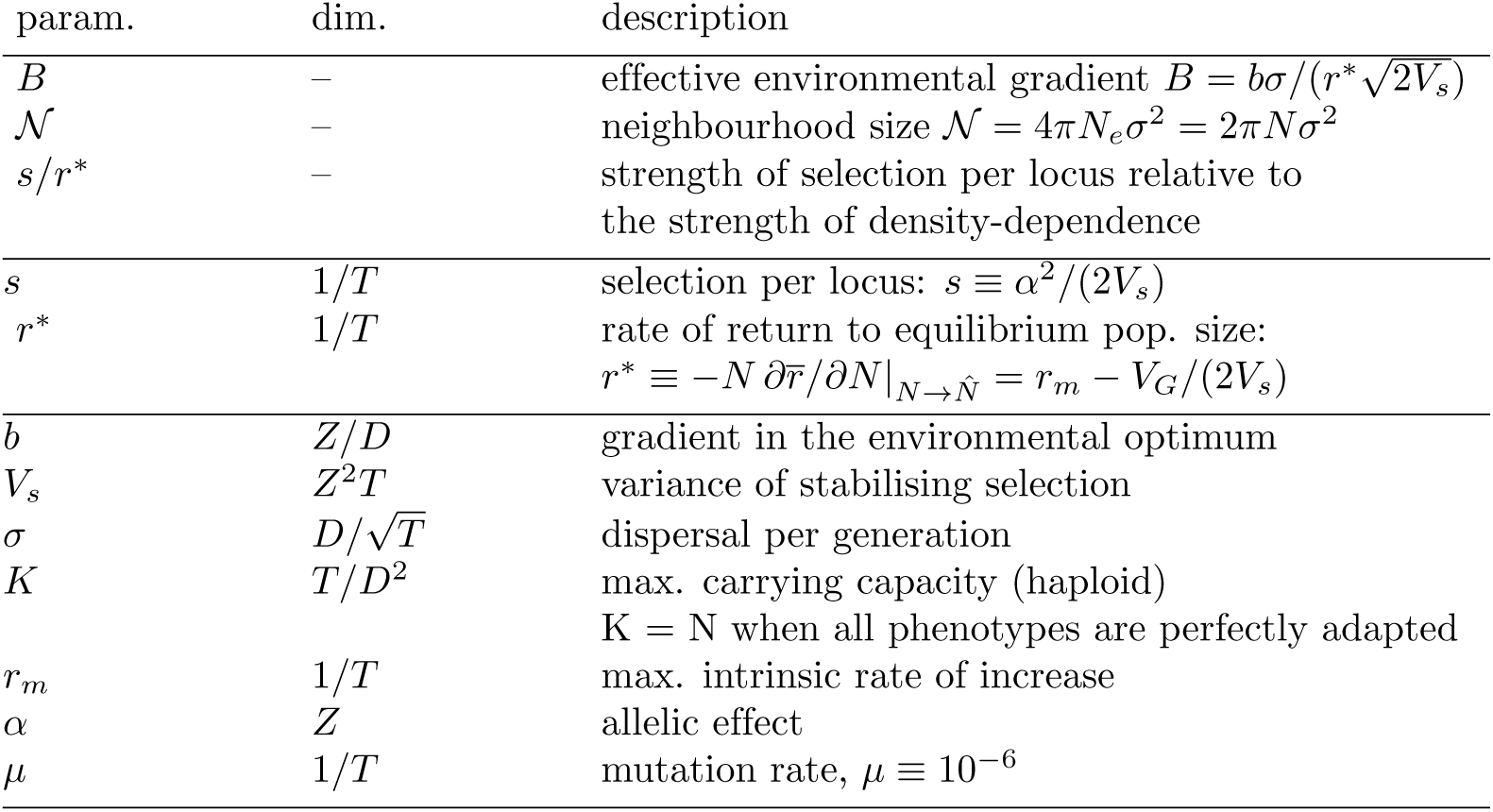
Three scale-free parameters: B, 𝒩 and *s/r*\* (top) describe the system. Middle section gives informative derived parameters. The bottom section gives seven parameters of the model before rescaling; where the seventh parameter, mutation rate *μ*, can be neglected because variance maintained by mutation-selection balance, *V_G,μ/s_* = 2*μV_s_n_l_*, is typically much smaller than variance generated by gene flow across environments, *V_G_* = *bσV_s_*. The middle column gives the dimensions of the parameters, where T stands for time, D for distance and Z for trait.

### Continuous model

For any given additive genetic variance *V_G_* (assuming a Gaussian distribution of breeding values), the change in the trait mean 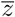 over time satisfies:

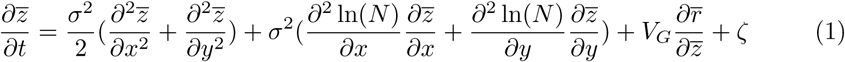

The first term gives the change in the trait mean due to migration with mean displacement of *σ*; the second term describes the effect of the asymmetric flow from areas of higher density. The third term gives the change due to selection, given by the product of genetic variance and gradient in mean fitness [58, Eq. 2]. The last term *ζ* gives the fluctuations in the trait variance due to genetic drift: 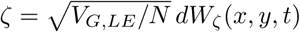, where *dW*_*_ represents white noise in space and time [34,59]. 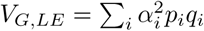 denotes genetic variance assuming linkage equilibrium.

The trait mean is 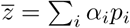 for a haploid model,where *p_i_* is the *i*-th allele frequency, *q_i_* = 1 − *p_i_* and *α_i_* is the effect of the allele on the trait – the change of the trait mean 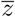 as frequency of locus *i* changes from 0 to 1. For both haploid and diploid models, the allele frequencies *p_i_* change as:

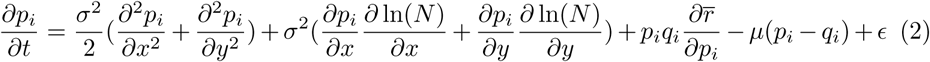

The expected change of allele frequency due to a gradient in fitness and local heterozygosity is 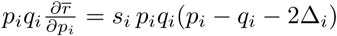, where selection at locus *i* is 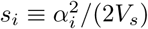 and 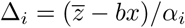 [13, Appendix 3]. Here, the fourth term describes the change due to (symmetric) mutation at rate *μ*. The last term *ϵ* describes genetic drift [34, Eq. 7]: 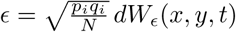; where *N* is the haploid population density.

Population dynamics reflect diffusive migration in two-dimensional habitat, growth due to the mean Malthusian fitness 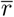, and stochastic fluctuations. The number of offspring follows a Poisson distribution with mean and variance of *N*, fluctuations in population numbers are given by [60]: 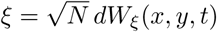:

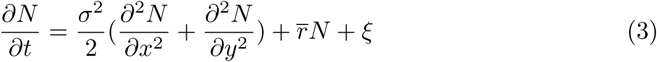

### Continuous model: Rescaling

The model can be simplified by rescaling [13, 58] time *t* relative to the strength of density dependence *r*\*, distance *x* relative to dispersal *σ*, trait *z* relative to strength of stabilising selection 1/(2*V_s_*) and local population size *N* relative to equilibrium population size with perfect adaptation: 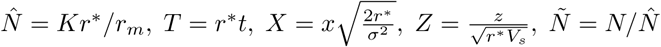. Note that near the equilibrium of a well-adapted population, 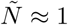.

The rescaled equations for evolution of allele frequencies and for demographic dynamics are:

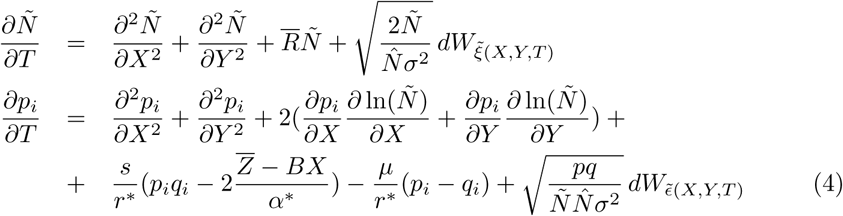

where 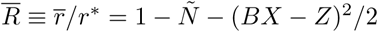.

The rescaled equations 4 and 5 show that four parameters fully describe the system. First, the effective environmental gradient, 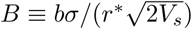. Second, the strength of genetic drift 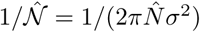. The parameter 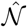 gives the neighbourhood size at an equilibrium with *uniform adaptation*. The third parameter is the strength of selection relative to the strength density dependence, *s/r*;* the scaled effect of a single substitution *α*\* also scales with *s/r\**: 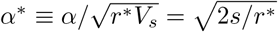. The effect of this third parameter *s/r*\* is expected to be small, because typically, *s* ⋘ *r*\*. Therefore assuming throughout that *s* is uniform across loci is a reasonably justified simplification. The fourth parameter, *μ*/*r*\*, will typically be very small, and will be neglected throughout. Table 1 (top) summarises the full set that describes the system.

## Supporting information

**S1 Fig. Adaptation to a steepening environmental gradient.** Shows the population trait mean, variance and population density in two-dimensional space.

**S2 Fig. Effect on genetic drift on a cline width in two-dimensional habitats.** The figure shows how the cline width – and hence the local genetic variance – decrease with genetic drift in two-dimensional habitats. In contrast to cline with in linear habitats [61], the effect of drift in nearly independent of selection.

**S3 Fig. Distribution of the parameters used in the randomised simulation runs from Fig. 2.**

## Acknowledgments

I am especially grateful to Nick Barton for his lasting knowledgeable advice and support. I would also like to thank my friends and colleagues for their suggestions and comments: Andrea Betancourt, David Field, Jarrod Hadfield, Joachim Hermisson, Sebastian Matuszewski (who also created the flower art), Glenn Marion, Richard Nichols, Tiago Paixão, Pavel Payne and Derek Setter. Last, I am would like to acknowledge the computer cluster team of IST Austria for their support long after I have the institute.

## Citations

1. Griggs RF. Observations on the behaviour of some species at the edges of their ranges. Bulletin of the Torrey Botanical Club. 1914;41:25–49.

2. Hargreaves AL, Samis KE, Eckert CG. Are species’ range limits simply niche limits writ large? A review of transplant experiments beyond the range. The American Naturalist. 2013;183(2):157–173.

3. Sexton J, McIntyre P, Angert A, Rice K. Evolution and ecology of species range limits. Annual Review of Ecology, Evolution, and Systematics. 2009;40:415–436.

4. Mayr E. Animal species and evolution. Harvard University Press; 1963.

5. Antonovics J. The nature of limits to natural selection. Annals of the Missouri Botanical Garden. 1976;63(2):224–247.

6. Gaston KJ. The structure and dynamics of geographic ranges. Oxford University Press; 2003.

7. Kawecki TJ. Adaptation to marginal habitats. Annual Review of Ecology, Evolution, and Systematics. 2008;39:321–342.

8. Eckert C, Samis K, Lougheed S. Genetic variation across species’ geographical ranges: the central–marginal hypothesis and beyond. Molecular ecology. 2008;17(5):1170–1188.

9. Cahill AE, Levinton JS. Genetic differentiation and reduced genetic diversity at the northern range edge of two species with different dispersal modes. Molecular ecology. 2016;25(2):515–526.

10. Takahashi Y, Suyama Y, Matsuki Y, Funayama R, Nakayama K, Kawata M. Lack of genetic variation prevents adaptation at the geographic range margin in a damselfly. Molecular ecology. 2016;25(18):4450–4460.

11. Pironon S, Papuga G, Villellas J, Angert AL, García MB, Thompson JD. Geographic variation in genetic and demographic performance: new insights from an old biogeographical paradigm. Biological Reviews. 2017;92(4):1877–1909.

12. Pujol B, Pannell JR. Reduced responses to selection after species range expansion. Science. 2008;321(5885):96–96.

13. Barton NH. Adaptation at the edge of a species’ range. In: Silvertown J, Antonovics J, editors. Integrating ecology and evolution in a spatial context. vol. 14. Blackwell; 2001. p. 365–392.

14. Haldane JBS. The relation between density regulation and natural selection. Proceedings of the Royal Society of London, B: Biological Sciences. 1956;145(920):306–308.

15. Nosil P, Crespi B. Does gene flow constrain adaptive divergence or vice versa? A test using ecomorphology and sexual isolation in *Timema cristinae* walking-sticks. Evolution. 2004;58(1):102–112.

16. Sexton JP, Strauss SY, Rice KJ. Gene flow increases fitness at the warm edge of a species’ range. Proceedings of the National Academy of Sciences. 2011;108(28):11704–11709.

17. Bourne EC, Bocedi G, Travis JM, Pakeman RJ, Brooker RW, Schiffers K. Between migration load and evolutionary rescue: dispersal, adaptation and the response of spatially structured populations to environmental change. Proceedings of the Royal Society of London B: Biological Sciences. 2014;281(1778):20132795.

18. Angert AL, Bayly M, Sheth SN, Paul JR. Testing range-limit hypotheses using range-wide habitat suitability and occupancy for the scarlet monkeyflower (*Erythranthe cardinalis*). The American Naturalist. 2018;191(3):E76–E89.

19. Gomulkiewicz R, Holt RD, Barfield M. The effects of density dependence and immigration on local adaptation and niche evolution in a black-hole sink environment. Theoretical Population Biology. 1999;55(3):283–296.

20. Holt RD, Gomulkiewicz R. How does immigration influence local adaptation? A reexamination of a familiar paradigm. The American Naturalist. 1997;149(3):563–572.

21. Alleaume-Benharira M, Pen I, Ronce O. Geographical patterns of adaptation within a species’ range: interactions between drift and gene flow. Journal of Evolutionary Biology. 2006;19(1):203–215.

22. Uecker H, Otto SP, Hermisson J. Evolutionary rescue in structured populations. The American Naturalist. 2014;183(1):E17–E35.

23. Barton N, Etheridge A. Establishment in a new habitat by polygenic adaptation. Theoretical Population Biology. 2018; p. 000–000. doi:10.1016/j.tpb.2017.11.007.

24. Kirkpatrick M, Barton N. Evolution of a species’ range. The American Naturalist. 1997;150(1):1–23.

25. Chevin L, Lande R. When do adaptive plasticity and genetic evolution prevent extinction of a density-regulated population? Evolution;. 2010;64(4):1143–1150.

26. Polechová J, Barton N. Limits to adaptation along environmental gradients. Proceedings of the National Academy of Sciences of the United States of America. 2015;112(20):6401–6406. doi:10.1073/pnas.1421515112.

27. Gilbert KJ, Whitlock MC. The genetics of adaptation to discrete heterogeneous environments: frequent mutation or large-effect alleles can allow range expansion. Journal of evolutionary biology. 2017;30(3):591–602.

28. Gilbert KJ, Sharp NP, Angert AL, Conte GL, Draghi JA, Guillaume F, et al. Local adaptation interacts with expansion load during range expansion: Maladaptation reduces expansion load. The American Naturalist. 2017;189(4):368–380.

29. Sæther BE, Lande R, Engen S, Weimerskirch H, Lillegård M, Altwegg R, et al. Generation time and temporal scaling of bird population dynamics. Nature. 2005;436(7047):99–102.

30. Polechová J, Barton N, Marion G. Species’ range: Adaptation in space and time. The American Naturalist. 2009;174(5):E186–204.

31. Christiansen FB. Hard and soft selection in a subdivided population. American Naturalist. 1975;109(965):11–16.

32. Wright S. Evolution in Mendelian populations. Genetics. 1931;16(2):97–159.

33. Kimura M, Weiss GH. The stepping stone model of population structure and the decrease of genetic correlation with distance. Genetics. 1964;49(4):561.

34. Barton NH, Depaulis F, Etheridge AM. Neutral evolution in spatially continuous populations. Theoretical Population Biology. 2002;61(1):31–48.

35. Wright S. Isolation by distance under diverse systems of mating. Genetics. 1946;31(1):39.

36. Slatkin M. Isolation by distance in equilibrium and non-equilibrium populations. Evolution. 1993;47(1):264–279.

37. Gould B, Moeller DA, Eckhart VM, Tiffin P, Fabio E, Geber MA. Local adaptation and range boundary formation in response to complex environmental gradients across the geographical range of *Clarkia xantiana ssp. xantiana*. Journal of Ecology. 2014;102(1):95–107.

38. Benning J, Eckhart VM, Geber MA, Moeller DA. Biotic interactions limit the geographic range of an annual plant: herbivory and phenology mediate fitness beyond a range margin. *submitted* 2018; p. -. doi:https://doi.org/10.1101/300590.

39. Case TJ, Taper ML. Interspecific competition, environmental gradients, gene flow, and the coevolution of species’ borders. The American Naturalist. 2000;155(5):583–605.

40. Spielman D, Brook BW, Frankham R. Most species are not driven to extinction before genetic factors impact them. Proceedings of the National Academy of Sciences of the United States of America. 2004;101(42):15261–15264.

41. Lande R. Genetics and demography in biological conservation. Science. 1988;241(4872):1455–1460.

42. Sagarin RD, Gaines SD. The ‘abundant centre’ distribution: to what extent is it a biogeographical rule? Ecology Letters. 2002;5(1):137–147.

43. Dallas T, Decker RR, Hastings A. Species are not most abundant in the centre of their geographic range or climatic niche. Ecology Letters. 2017;20(12):1526–1533. doi:10.1111/ele.12860.

44. Cotto O, Wessely J, Georges D, Klonner G, Schmid M, Dullinger S, et al. A dynamic eco-evolutionary model predicts slow response of alpine plants to climate warming. Nature Communications. 2017;8.

45. O’Brien EK, Higgie M, Reynolds A, Hoffmann AA, Bridle JR. Testing for local adaptation and evolutionary potential along altitudinal gradients in rainforest Drosophila: beyond laboratory estimates. Global change biology. 2017;23(5):1847–1860.

46. Slatkin M, Barton NH. A comparison of three indirect methods for estimating average levels of gene flow. Evolution. 1989;43(7):1349–1368.

47. Rousset F. Genetic differentiation and estimation of gene flow from F-statistics under isolation by distance. Genetics. 1997;145(4):1219–1228.

48. Charlesworth B, Charlesworth D, Barton NH. The effects of genetic and geographic structure on neutral variation. Annual Review of Ecology, Evolution, and Systematics. 2003;34(1):99–125.

49. Whitlock MC, McCauley DE. Indirect measures of gene flow and migration: *F_ST_* = 1/(4N*m* + 1). Heredity. 1999;82(2):117–125.

50. Blows M. A tale of two matrices: multivariate approaches in evolutionary biology. Journal of evolutionary biology. 2007;20(1):1–8.

51. Agrawal AF, Stinchcombe JR. How much do genetic covariances alter the rate of adaptation? Proceedings of the Royal Society of London B: Biological Sciences. 2009;276(1659):1183–1191.

52. Colautti RI, Eckert CG, Barrett SC. Evolutionary constraints on adaptive evolution during range expansion in an invasive plant. Proceedings of the Royal Society B: Biological Sciences. 2010;277(1689):1799–1806.

53. Duputié A, Massol F, Chuine I, Kirkpatrick M, Ronce O. How do genetic correlations affect species range shifts in a changing environment? Ecology letters. 2012;15(3):251–259.

54. Hine E, McGuigan K, Blows MW. Evolutionary constraints in high-dimensional trait sets. The American Naturalist. 2014;184(1):119–131.

55. Lande R. Natural selection and random genetic drift in phenotypic evolution. Evolution. 1976;30(2):314–334.

56. Barton N. How does epistasis influence the response to selection? Heredity. 2017;118(1):96–109. doi:10.1038/hdy.2016.109.

57. Polechová J, Barton NH. Speciation through competition: A critical review. Evolution. 2005;59(6):1194–1210.

58. Pease CM, Lande R, Bull JJ. A model of population growth, dispersal and evolution in a changing environment. Ecology. 1989;70(6):1657–1664.

59. Nagylaki T. Random genetic drift in a cline. Proceedings of the National Academy of Sciences of the United States of America. 1978;75(1):423–426.

60. Lande R, Engen S, Sæther BE. Stochastic population dynamics in ecology and conservation. Oxford University Press; 2003.

61. Polechová J, Barton N. Genetic drift widens the expected cline but narrows the expected cline width. Genetics. 2011;1(189):227–235.

